# Amplitude modulation encoding in auditory cortex: Comparisons between the primary and middle lateral belt regions

**DOI:** 10.1101/2020.03.05.979575

**Authors:** Jeffrey S. Johnson, Mamiko Niwa, Kevin N. O’Connor, Mitchell L. Sutter

## Abstract

In macaques, the middle lateral auditory cortex (ML) is a belt region adjacent to primary auditory cortex (A1) and believed to be at a hierarchically higher level. Although ML single-unit responses have been studied for several auditory stimuli, the ability of ML cells to encode amplitude modulation (AM) – an ability which has been widely studied in A1 – has not yet been characterized. Here we compare the responses of A1 and ML neurons to amplitude modulated (AM) noise in awake macaques. While several of the basic properties of A1 and ML responses to AM noise are similar, we found several key differences. ML neurons do not phase lock as strongly, are less likely to phase lock, and are more likely to respond in a non-synchronized fashion than A1 cells, consistent with a temporal-to-rate transformation as information ascends the auditory hierarchy. ML neurons tend to have lower temporally (phase-locking) based best modulation frequencies than A1. At the level of ML, neurons that decrease firing rate with increasing modulation depth become more common than in A1. In both A1 and ML we find a prevalent class of neurons with excitatory rate responses at lower modulation frequencies and suppressed rate responses relative to the unmodulated carrier at middle modulation frequencies.

## INTRODUCTION

The middle lateral belt area of auditory cortex (ML) is a region in the auditory belt adjacent to primary auditory cortex (A1), and is considered to sit at a higher level of the auditory hierarchy due to its lack of input from the medial geniculate nucleus of the thalamus (Kaas and Hackett 2000). Given its location on the cortical sheet as well as its connections (Romanski et al. 1999), latency (Camalier et al. 2012), and tuning properties to auditory objects and spatial location (Rauschecker and Tian 2000; Tian et al. 2001; Woods et al. 2006) ML appears to straddle the proposed “what” and “where” pathways in auditory cortex. Although it is tonotopic like A1 (Rauschecker et al. 1995) it prefers more complex stimuli such as bandpass noise (Rauschecker et al. 1995; Rauschecker and Tian 2004) and frequency-modulated sweeps (Tian and Rauschecker 2004) to tones.

Although there is a rudimentary knowledge of ML response properties, a detailed examination of many basic auditory stimuli is still lacking. In particular, ML responses to amplitude modulated stimuli remain largely uncharacterized. Amplitude modulation (AM) is a common feature of sound and an important information containing parameter for natural sounds, including animal vocalizations and speech (e.g. Rosen 1992; Drullman et al. 1994; Shannon et al. 1995; Liu et al. 2003; Singh and Theunissen 2003; Zeng et al. 2005; Jin and Nelson 2006; Narayan et al. 2006; Cohen et al. 2007; Elliot and Theunissen 2009; Geffen et al. 2011; Xiang et al. 2013; Gervain and Geffen 2019) as well as a feature useful for segregating and attending to sounds, more commonly referred to as the cocktail party problem (Bregman 1990; Yost 1991; Grimault et al. 2002; Steinschneider et al. 2003; Itatani and Klump 2009; Bohlen et al. 2014; Hershenhoren and Nelken 2017; Yamagishi et al. 2017) Despite its importance in auditory scene analysis, we know very little about how higher areas of cortex process AM.

Because of their periodic nature, sine-AM stimuli are one of the simplest tools for investigating the temporal aspects of auditory processing. AM stimuli have been used to characterize every stage of the auditory system from auditory nerve to A1 in various animal models (reviewed in Joris et al. 2004). A1 in the non-human primate has been particularly well-studied with AM (e.g. Lu et al. 2001; Liang et al. 2002; Bendor and Wang 2007; Malone et al. 2007; Bendor and Wang 2008; Scott et al. 2011; Yin et al. 2011; Johnson et al. 2012; Malone et al. 2013; Bohlen et al. 2014; Overton and Recanzone 2016; Hoglen et al. 2018), establishing a good opportunity for the characterization of ML in comparison to A1. Recent reports have pointed to differences in the processing of AM noise stimuli in A1 and ML with regard to higher level processes such as task engagement (Niwa et al. 2015) and decision making (Niwa et al. 2013) but there remains a gap in our knowledge of simpler, more fundamental response properties to AM stimuli in ML.

One consistent finding in studies of neural responses to AM stimuli is that there is a temporal-to-rate transformation as information ascends the auditory hierarchy. High frequency phase-locking cutoffs gradually decrease and a class of cells that codes for modulation frequency with spike rate but does not phase lock to the stimulus itself emerges (e.g. Schreiner and Urbas 1988; Lu et al. 2001; Lu and Wang 2004; Bendor and Wang 2007; Gao and Wehr 2015). Because ML is putatively at a higher level than A1 in the auditory hierarchy, a continuation of this temporal-to-rate transformation from A1 to ML would be consistent with the hierarchical placement of ML.

In this study we compare various aspects of AM noise processing in A1 and ML of the awake non-behaving macaque. We find much in common between the two areas, but the results indicate a shift away from temporal coding at the level of ML, consistent with an ascending temporal-to-rate transformation. The results also suggest that the gradual emergence of a dual code – wherein modulation may be encoded with either an increase or a decrease in firing rate – previously reported in behaving macaques (Niwa et al. 2013) also occurs in non-behaving macaques. Additionally, in both A1 and ML we find an abundant class of cells not widely reported in cortex that shows excitatory rate responses at lower modulation frequencies and suppressed rate responses at middle modulation frequencies relative to the unmodulated carrier – we call these cells excitation-suppression (E/S) cells. These cells have a low-frequency region that is typically phase-locked, and a high-frequency region that is typically not phase-locked and has firing rates similar to those seen to the unmodulated carrier, suggesting that the high frequency region may encode the carrier rather than the modulation.

## MATERIALS AND METHODS

### Subjects

In this study data were collected from the right hemispheres of three adult rhesus macaque monkeys, two females (monkeys V and W) and one male (monkey X). A1 single neuron recordings were collected in all three monkeys. ML single neuron recordings were collected in monkeys W and X. All procedures conformed to the United States Public Health Service (PHS) policy on experimental animal care and were approved by the UC Davis animal care and use committee.

### Stimulus generation and presentation

Acoustic stimuli were 800-ms sinusoidally amplitude-modulated (AM) broadband noise bursts. AM stimuli had modulation frequencies (MFs) of 2.5, 5, 10, 15, 20, 30, 60, 120, 250, 500, and 1000 Hz, and were at 100% modulation depth. In addition an unmodulated broadband noise burst was used. All stimuli were created from the same random number sequence (“frozen” noise) and were thus identical prior to the application of the modulation envelope.

The digital version of each stimulus was created in Matlab (MathWorks) and was given a 5-ms cosine ramp at onset and offset. Digital-to-analog conversion was performed at a sampling rate of 100 kHz with a D/A converter (Power1401; Cambridge Electronic Design). The resulting analog signal was passed through both a programmable attenuator (PA5; Tucker-Davis Technologies) and a passive attenuator (LAT-45; Leader) before being amplified (MPA-200; Radio Shack) and output to a speaker inside a sound booth. We used two different sound booths (both manufactured by IAC) with different speakers. One booth measured 2.9 × 3.2 × 2.0 m and had a Radio Shack PA-110 speaker positioned 1.5 m in front of the subject; the other booth measured 1.2 × 0.9 × 2.0 m and had a Radio Shack Optimus Pro-7AV speaker positioned 0.8 m in front of the subject. In both cases the speaker was positioned at ear level. Stimulus intensity was calibrated with a sound level meter (model 2231; *Brüel & Kjær*) to 63 dB sound pressure level at the position of the subjects’ pinnae.

### Behavioral procedure

During the period of this study, the animals were maintained on fluid regulation. For the recording sessions used here, the animals alternated between passive blocks (no task) with alertness maintained by occasional liquid reward and active blocks where the animals were given liquid rewards for correctly discriminating AM stimuli from unmodulated stimuli. Only data from passive blocks are used in this study; some results from active block data have been reported elsewhere (Niwa et al. 2012; Niwa et al. 2013; Niwa et al. 2015).

### Physiological recording

Each animal was chronically implanted with a titanium head post and a CILUX recording chamber (Crist Instruments) which was located over parietal cortex for access to A1 and ML. A plastic grid with 27-gauge holes arranged over a 15 × 15 mm area in 1 mm intervals was mounted on the recording chamber. For each recording a stainless steel transdural guide tube was inserted through the grid. A high impedance tungsten microelectrode (1–4 MΩ, FHC; 0.5–1 MΩ, Alpha-Omega) was inserted into the guide tube and lowered through parietal cortex into A1 or ML with a hydraulic microdrive (FHC). Recordings were made while the animal was head-restrained by the head post in an “acoustically transparent” custom-made primate chair made of wire.

Electrophysiological signals were passed through an amplifier (A-M Systems model 1800) and filtered (0.3-10kHz; A-M Systems model 1800 and Krohn-Hite model 3382) before being sent to an A/D converter (Power 1401; Cambridge Electronic Design), sampled at a rate of 50 kHz and saved to hard disk. Action potentials were spike-sorted offline using a waveform-matching algorithm (Spike2; Cambridge Electronic Design).

### Data analysis

All data analysis was performed using Matlab (MathWorks). All statistical comparisons between counts or percentages were performed with a z-test for independent proportions.

### Measures of phase locking

#### Phase-projected vector strength

We used phase-projected vector strength (*VS*_*PP*_) as our primary measure of phase locking. Conceptually, *VS*_*PP*_ is a trial-based measure of synchrony which penalizes trial-based vector strength (*VS*) values for having a mean phase which is misaligned with the cell’s overall mean phase (Yin et al. 2011). The standard formula for vector strength is

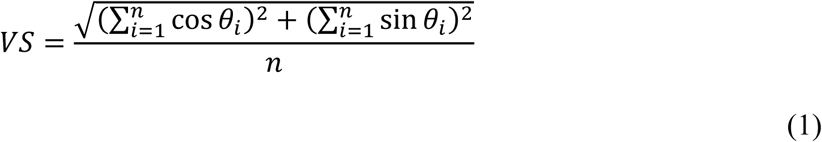

where *VS* is the vector strength, n is the number of spikes, and *θ*_i_ is the phase of each spike in radians, calculated by

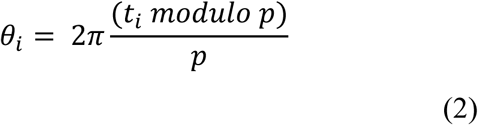

where *t*_i_ is the time of the spike in ms relative to the onset of the stimulus and *p* is the modulation period of the stimulus in ms (Goldberg and Brown 1969; Mardia and Jupp 2000).

Phase-projected vector strength (*VS*_*PP*_) was calculated on a trial-by-trial basis as follows:

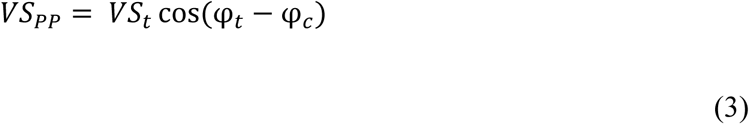

where *VS*_*PP*_ is the phase-projected vector strength per trial, *VS*_*t*_ is the vector strength per trial, calculated as in Eq. 1, and φ_t_ and φ_c_ are the trial-by-trial and mean phase angle in radians respectively, calculated for each stimulus condition

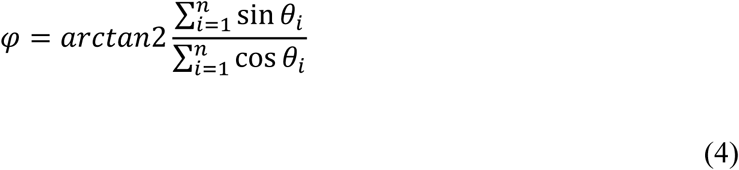

where *n* is the number of spikes per trial (for φ_t_) or across all trials (for φ_c_) and arctan2 is a modified version of the arctangent that determines the correct quadrant of the output based on the signs of the sine and cosine inputs (Matlab, ***atan2***). A cell that fired no spikes was assigned a *VS*_*PP*_ of zero.

We also calculated cycle-by-cycle vector strength (*VS*_*CC*_) to estimate the reliability of phase locking (Yin et al. 2011). *VS*_*CC*_ combines measures of how precise the timing of a neuron’s firing is and how often a neuron successfully responds to an AM cycle. To calculate *VS*_*CC*_, one *VS*_*PP*_ value is calculated for each complete AM cycle of the stimulus and then all these values are averaged together to give *VS*_*CC*_ for a given trial. If no spike is fired for a given cycle, a value of 0 is used for that cycle. As such *VS*_*CC*_ is a measure that incorporates reliability of the phase locking on a cycle by cycle basis. A cell that fires at exactly the same time within a cycle, but fails to fire on half of the cycles will have a *VS*_*PP*_ of 1.0, but a *VS*_*CC*_ of 0.5.

### Determining significance of phase locking and firing rate

To determine if a cell was responsive to AM via either firing rate or phase locking we performed a statistical analysis. Significance of phase locking was calculated for each cell at each recorded modulation frequency (MF) with a two-tailed t-test (P < 0.05, corrected for 11 multiple comparisons) evaluating the null hypothesis that the distribution of *VS*_*PP*_ values in response to AM was not different than the distribution of *VS*_*PP*_ values in response to unmodulated noise. Rate significance was calculated in the same fashion, but using the distributions of firing rates. A cell was considered to “detect” AM if there was significance for at least one MF. If a neuron failed this test of whether it could detect AM it was classified as non-responsive (NR). Each neuron was evaluated separately for AM responsiveness based on phase locking or firing rate. For any measure for which the neuron was responsive, a curve fit (see below) was performed on the modulation transfer function (MTF). If the neuron was NR for a measure the MTF was not fit for that measure. Analogous tests were performed for unmodulated noise detection against a 200-ms period of spontaneous activity preceding each stimulus.

### Fitting temporal modulation transfer functions (tMTFs)

For neurons whose VS_PP_-based phase locking was not classified as NR, curve fits were used to place the tMTF into one of four categories: low pass (LP), bandpass (BP), high pass (HP), or No Fit. To achieve this, responsive tMTFs were separately fitted (Matlab “fmincon” function) with three functions; a (logistic) sigmoid (Eq. 5), a Gaussian (Eq. 6), and a log-transformed Gaussian function (Eq. 7). All three functions have four free parameters determining the y-offset (*a*), the height (*b*), the x-center (*µ*), and the slope (*s*). Because they had the same number of free parameters, error minimization (described below after the equation constraints) could be used to determine which function (if any) was the best fit.

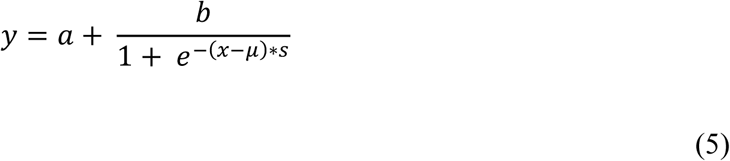

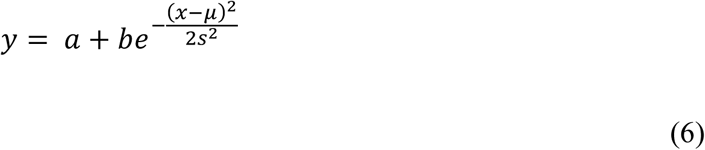

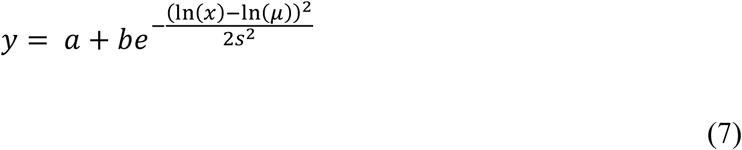

The constraints on the parameters were set as follows:

For all fits: 0.9 × (min of data) ≤ *a* ≤ 1.1 × (min of data); 0 ≤ *b* ≤ 1.03 × (max-min of data); 0 ≤ *µ* ≤ 2000 Hz; The constraint on the height parameter prevented excessively high amplitude extrapolations.

Logistic: −1 ≤ *s* ≤ 1. The slope factor restricted the 90% width of the logistic to be about 6 Hz or greater. A negative slope parameter allows the logistic to fit a low pass MTF while a positive slope parameter allows it to fit a high pass MTF.

Gaussian: 3 ≤ *s* ≤ 213. The slope factor restricted the full width at half height (FWHH) of the Gaussian fit to lie between about 7 and 500 Hz.

Log-Transformed Gaussian: 0.05 ≤ *s* ≤ 2.36. The slope factor restricted the FWHH of the log-transformed Gaussian fit to a minimum of about 1/6 of an octave and a maximum of about 8 octaves.

### Determining best fit function and classifying tMTFs

A minimization was used to determine which of the three functions provided the best fit to the tMTF. The minimization in the tMTF fit procedures was weighted by the synchronized spike count (*SSC*, = *VS* × spike count) at each MF to emphasize fitting to the more reliable points in the tMTF. The coefficient of determination (*CoD*) was calculated for each fit as follows:

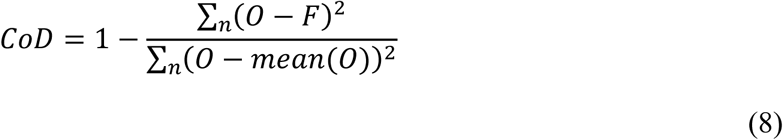

where *O* is the set of *n* observed points in the MTF and *F* is the corresponding set of *n* points from the resulting fitted function. The *CoD* is bounded by 1 on the high end, which indicates that all fitted points match the observed points exactly, but is unbounded on the low end. A *CoD* of zero indicates that the fit is equally good to a horizontal line plotted through the mean of the observed values. tMTFs for which none of the three fits (logistic, Gaussian, log-transformed Gaussian) had a *CoD* greater than 0.5 were categorized as No Fit (note that this is different than non-responsive, NR, because there were significant responses or phase locking to AM). tMTFs with at least one *CoD* greater than 0.5 which were best fit (i.e. had the highest *CoD*) by a sigmoid function were categorized as LP or HP based on the direction of the sigmoid’s inflection (in our data set we did not observe any HP tMTFs). tMTFs that were best fit by one of the Gaussian functions were categorized as BP, unless the *VS*_*PP*_ value at the lowest MF (or highest MF) was at least 90% of the highest *VS*_*PP*_ value found in that neuron, in which case the tMTF was categorized as LP (or HP) for lack of sufficient evidence of a downturn.

### Classifying rate-based modulation transfer function (rMTFs)

The same categorization that was done on tMTFs was initially performed on rMTFs. However, we observed that a large number of our rMTFs did not fit neatly into the categories defined for tMTFs above. Specifically, many had multiple peaks or dips, so we modified the fitting procedure in order to quantify this.

Therefore, all rMTFs were also fit in a separate multi-step procedure with a two-curve fit. The first step of the two-curve fit was to fit the rMTF with the best of a Gaussian or log-transformed Gaussian, with the minimization weighted by the synchronized spike count *(SSC)*, which emphasizes fitting synchronized and/or higher spike count regions of the MTF. This first fit was constrained to have a floor at the spontaneous firing rate of the cell. The result of this fit was subtracted from the rMTF to obtain a residual MTF, and the residual MTF was subsequently fit with the best of a Gaussian or log-transformed Gaussian, with the minimization weighted by the quantity max(*SSC*) – *SSC*_*n*_ at each MF, so the weighting is higher for points (n in equation) with less synchronization. (Max(SSC) is the point on the MTF with the maximum synchronized spike count.) This second fit was additionally constrained to have a floor of zero. These two fits were added together to obtain the two-curve fit.

The two-curve fit aided identification of a commonly observed rMTF shape that we called excitation/suppression (E/S) rMTFs. E/S have excitatory rate responses at lower MFs and suppressed rate responses relative to the unmodulated carrier at middle MFs, and return to baseline (firing rate to unmodulated noise) at higher MFs.

As with tMTFs, rMTFs without any responses significantly different from the response to unmodulated noise at any modulation frequency were categorized as NR, and rMTFs for which none of the four fits (logistic, Gaussian, log-transformed Gaussian, two-curve) had a *CoD* greater than 0.5 were categorized as No Fit. rMTFs that were best fit (i.e. had the highest *CoD*) by a function other than the two-curve fit were categorized using the same method as tMTFs; only rMTFs that were best fit by the two-curve fit were considered potential E/S rMTFs. Potential E/S rMTFs were subjected to three tests to discourage “overfitting” with our two-curve fit when a one-curve fit would have been sufficient. First, we required that the two-curve fit clearly be the best fit. Our criterion here was that the *CoD* of the two-curve fit must be at least 0.02 greater than the *CoD* of the best one-curve fit. Second, we required that the two-curve fit consist of largely non-overlapping curves. Here, we defined the half-height as the cutoff for each curve. We required that that these cutoffs not overlap in MF, and additionally that at least one collected MF data point fell between the two cutoffs (i.e., high cutoff of curve 1 < MF of a collected data point < low cutoff of curve 2). Third, we required that neither curve could be more than four times the height of the other, which prevents solutions that overfit a noisy tail of a one-curve solution. MTFs that met all three of these criteria were considered E/S, otherwise they were classified based on their best one-curve fit.

### Categorizing whether neurons increase, or decrease activity in response to modulation

We defined increasing and decreasing cells somewhat differently than in Niwa et al. 2013 because whether a cell increases or decreases its response to modulation may depend on the tested MF. For each MF in the MTF, we compared the trial-by-trial spike count or *VS*_*PP*_ for the 100% depth stimulus against the unmodulated stimulus to determine if it was greater than or less than unmodulated noise (one-sided t-test for each comparison, corrected for 11 MFs). Cells with at least one MF significantly greater than unmodulated and at least one MF significantly less than unmodulated were classified as “mixed”, otherwise a cell was classified as “increasing” if at least one MF was significantly greater than unmodulated or “decreasing” if at least one MF was significantly less than unmodulated.

### Cutoff frequencies and bandwidths

For logistic fits, the high-frequency cutoff was calculated as the half-height point of the curve. For both Gaussian fits, low-frequency and high-frequency cutoffs were selected as the two half-height points on the curve, and the bandwidth was calculated as the full width at half height (FWHH). Low-frequency cutoff and bandwidth values were rejected for any Gaussian fit where the low-frequency cutoff was less than zero. In the two-curve fits, cutoffs and bandwidths were calculated for each curve individually.

## RESULTS

We recorded spiking activity from 250 single neurons in primary auditory cortex (A1) of three macaques (monkey V, 21 units; monkey W, 120 units; monkey X, 109 units) and from 138 single neurons in the middle lateral auditory cortex (ML) of two macaques (monkey W, 68 units; monkey X, 70 units) while the animals were awake and presented with unmodulated noise and 100% amplitude modulated (AM) noise.

### Modulation transfer functions

Responses to AM for four ML neurons exemplify their heterogeneity (Fig. 1). Figure 1A depicts a neuron that phase locks to the AM stimulus with a high firing rate up to 30 Hz and then continues to phase lock well with a lower cycle-by-cycle reliability up to 250 Hz. Figure 1B depicts another phase-locking cell that typically fires only one spike per AM cycle. Figures 1C and 1D show two neurons that only produce non-synchronized responses to AM-noise, one with a narrow rate-based modulation transfer function (rMTF) and the other with a much broader rMTF. We attempted to categorize all of our MTFs by shape into various classes, including the familiar low pass, bandpass and high pass classes, using an automated fitting procedure (see Methods). Cells which had a significant response for at least one modulation frequency (MF) but which did not achieve a significant fit for any MTF class were categorized as “no fit”. Unlike temporal-based MTFs (tMTFs), rMTFs (including the one in Fig. 1B) often best fit into a novel class - the excitation-suppression (E/S) class, which is described in more depth below.

**Figure 1:**
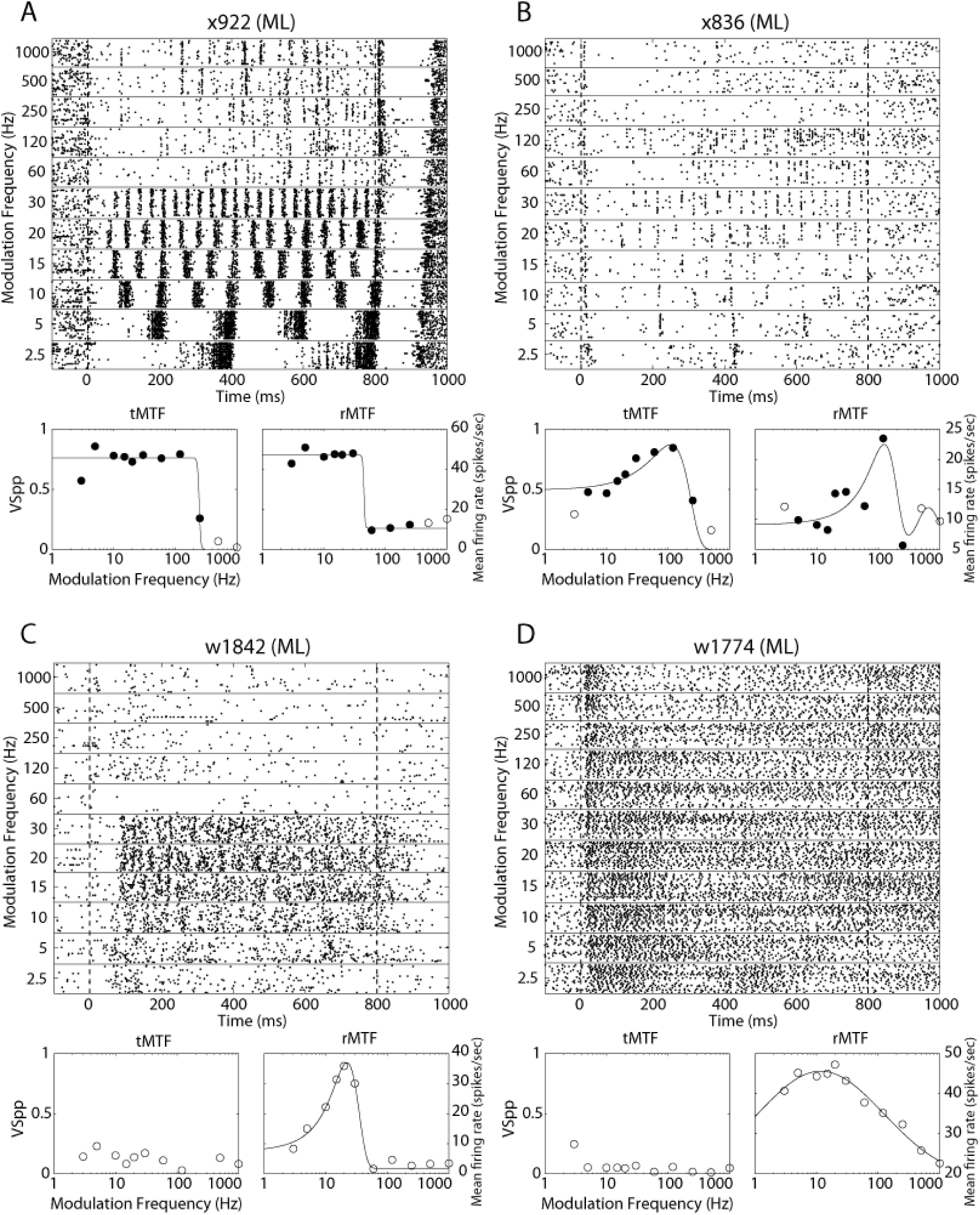
Example responses of ML neurons to AM noise. The responses to AM noise, and the modulation transfer functions are shown for four different example neurons (Panels A-D). For each neuron we show the responses to the eleven tested modulation frequencies with raster plots on the top of the panel. Below each raster plot are temporal (left, tMTF) and rate (right, rMTF) modulation transfer functions (MTFs) for the same neurons. Filled dots on the MTFs indicate modulation frequencies at which there is significant phase locking (t-test, P < 0.05 corrected for 11 comparisons) - therefore, the same modulation frequencies on both temporal and rate MTFs are filled.

There were some differences in MTF and response properties between ML and A1 neuronal populations (Table 1). The proportion of cells that phase lock to AM-noise is lower in ML than in A1 (172/250, 69% in A1; 66/138, 48% in ML; P = 5×10^−5^, all statistical comparisons between counts or percentages are performed with a z-test for independent proportions). Among cells that phase lock to AM, high pass tMTFs are not found in A1 or ML and bandpass is the most common tMTF type in both areas. However, there is a much higher percentage (27%) of neurons with low pass tMTFs in ML than in A1 (7%; P = 21×10^−5^) and concomitantly more bandpass tMTFs in A1 than in ML (P = 1×10^−5^). All of these observations support a reduction in phase-locking ability in ML consistent with a temporal-to-rate transformation from A1 to ML.

**Table 1:** MTF Classification: Values on the left are in cell count, values on the right are in percentage. Denominator for “Not Responsive” cells is total cells recorded. Denominator for all others is total responsive cells (or for “synch” columns, total responsive cells within the given MTF class). Neurons are considered to synchronize if they have at least one modulation frequency with significant phase locking whose firing rate was at or above the half-height of the fitted Gaussian/sigmoid function for the MTF. Neurons that are not responsive to rate do not synchronize by our definition, even if they exhibit some significant phase locking.

|  | Temporal |  | Rate |  |  |  |
| --- | --- | --- | --- | --- | --- | --- |
|  | A1 | ML | A1 |  | ML |  |
|  |  |  | Total | % that Synch | Total | % that Synch |
| Not Responsive | 78/250 31.2% | 72/138 52.2% | 72/250 28.8% | 0/72 0% | 50/138 36.2% | 0/50 0% |
| No Fit | 24/172 14.0% | 15/66 22.7% | 23/178 12.9% | 1/23 4.3% | 23/88 26.1% | 1/23 4.3% |
| Low Pass | 12/172 7.0% | 18/66 27.3% | 3/178 1.7% | 1/3 33.3% | 5/88 5.7% | 4/5 80.0% |
| Bandpass | 136/172 79.1% | 33/66 50.0% | 71/178 39.9% | 43/71 60.6% | 21/88 23.9% | 10/21 47.6% |
| High Pass | 0/172 0% | 0/66 0% | 8/178 4.5% | 0/8 0% | 5/88 5.7% | 1/5 20.0% |
| Excitation/<br>Suppression | N/A | N/A | 73/178 41.0% | Low: 58/73 79.5%<br>High: 2/73 2.7% | 34/88 38.6% | Low: 29/34 85.3%<br>High: 2/34 5.9% |

The proportion of cells that change their firing rate to AM is not significantly different between the two areas (178/250, 71% in A1; 88/138, 64% in ML; P = 0.13). For rMTF shape classification, there are only a few salient differences between the A1 and ML populations. For one, ML shows an increase in the number of responsive cells that were not well fit (P = 71×10^−3^), which could reflect an increase in the complexity of MTFs in higher levels of the auditory system. There are also fewer bandpass rMTFs in ML than A1 (P = 0.01) which seems to roughly correspond to the increase in no-fit rMTFs. We performed a separate analysis of rate-responsive cells that also synchronized to AM. Here, rate-responsive cells are considered “synchronized” if they also exhibit significant phase locking for at least one MF within the cutoffs of the rMTF fit (defined as the MFs corresponding to the half-max of the fit). Between A1 and ML, there is no significant difference in the proportion of synchronized cells for any rMTF class. Bandpass, low pass and excitation-suppression (E/S) rMTFs are the only classes that commonly synchronize.

### Encoding of AM versus responding to the carrier

Modulation transfer functions describe, in terms of general shape, how a neuron responds to sounds with various modulation frequencies, but they do not inherently speak to a neuron’s ability to detect a stimulus or a stimulus feature. Spiking activity can be used to detect the presence of a stimulus or stimulus feature only if it differs from spiking activity when the stimulus or feature is not present. When neurons are presented with AM sounds, we can define two kinds of detection. First, the neuron can detect the presence of a sound as compared to the absence of a sound – this is stimulus detection, and the appropriate comparison is with the neuron’s spontaneous firing. Alternatively, the neuron can detect the presence of AM in a sound as compared to the absence of AM in a sound – this is modulation detection, and the appropriate comparison is with the neuron’s response to an unmodulated stimulus (carrier).

Figure 2 demonstrates these detection differences in A1 and ML. The left panel depicts the percentage of cells exhibiting stimulus detection, while the right panel shows the extent of AM detection. ML exhibits a decrease in the ability to use synchronized responses (*VS*_*PP*_ + *SC*) for detection, but an increase in the ability to use exclusively non-synchronized responses (*SC* Only) – those that exhibit a change in firing rate without phase locking to the stimulus. This shift from temporal coding to rate coding in ML is consistent with a move up in the auditory hierarchy. In addition, there is an increase in the percentage of cells that do not detect the stimulus in ML compared to A1, but the percentage of cells that detect modulation is not significantly different between the two areas. There is no change across area in the percentage of exclusively synchronized cells (*VS*_*PP*_ Only), and these cells only account for 10 percent or less of the population.

**Figure 2:**
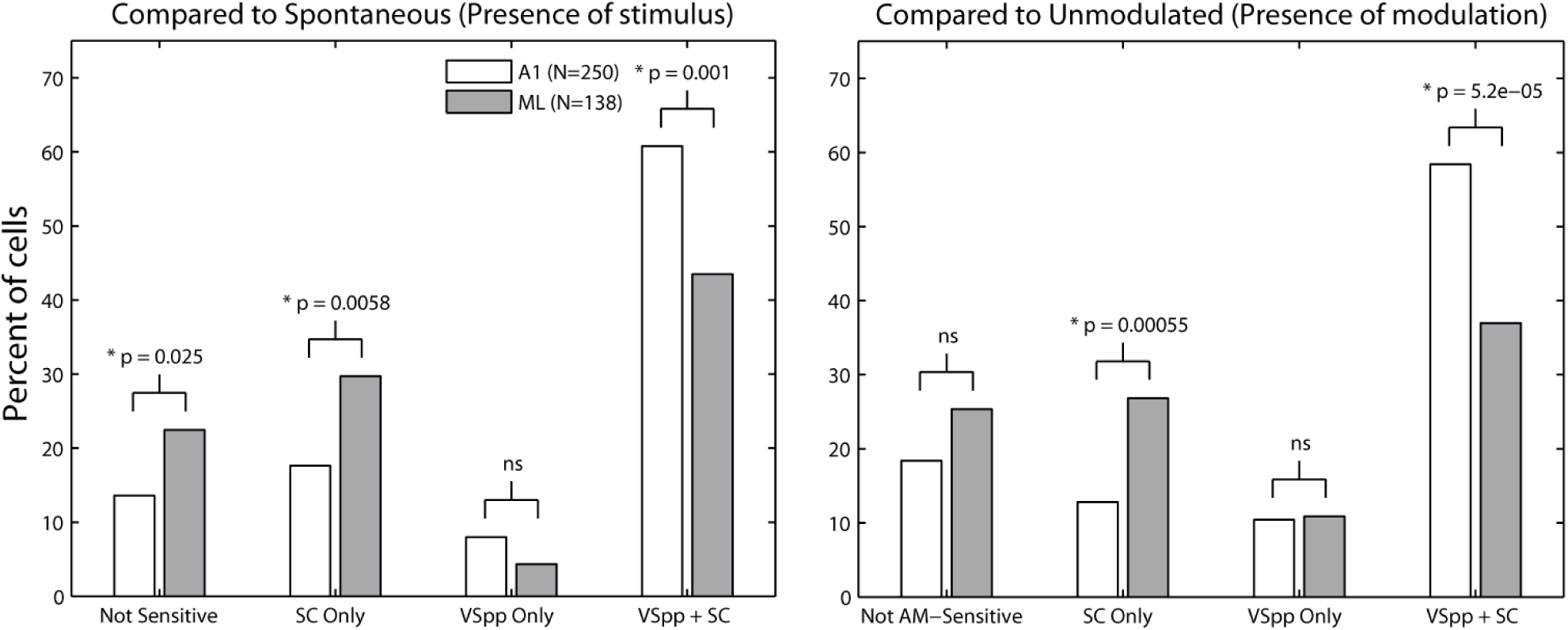
Detection of a stimulus and detection of modulation. Left, percent of cells in A1 (open) and ML (shaded) that detect the stimulus (compared against spontaneous firing). Cells that failed to detect the presence of a stimulus with either spike count or vector strength are labelled as Not Sensitive. Cells that could detect the presence of a stimulus (relative to spontaneous) with spike counts, but not with vector strength are labelled SC Only. Cells that could detect the presence of a stimulus (relative to spontaneous activity) with vector strength, but not with spike counts are labelled VS_PP_ Only. Cells that showed significant differences from spontaneous for both spike counts and vector strength are labelled as VS_PP_ + SC. For each category an A1/ML comparison was made using a z-test for the difference of independent proportions (P-value noted above bars). Right, same as left, except the comparison was made against the unmodulated stimulus and represents detection of amplitude modulation at 100% depth.

The general pattern for both detection of a stimulus and detection of AM is very similar. The only statistically significant difference in detection in either area is an increase in ML of exclusively-synchronized cells (*VS*_*PP*_ Only) detecting AM relative to those detecting the stimulus (P = 0.04). However, the *VS*_*PP*_ Only cells were the smallest class (4% of cells detecting the stimulus and 11% of cells detecting modulation), and therefore whether this difference is important is hard to determine.

### Best modulation frequencies

The top and middle panels of Figure 3 show that the distribution of best modulation frequencies (BMFs) for both rate (rBMF, spike count) and temporal (tBMF, *VS*_*PP*_) measures can be greatly influenced by the range of modulation frequencies tested. The top panel depicts the A1 BMF count in the present data looking only at the frequencies which were tested in a previous report of A1 BMFs (Yin et al. 2011). The distribution of tBMFs is similar to that found previously. The distribution of rBMFs is a bit different – in the current sample about half of the cells would have shown an rBMF of 5 Hz if tested only at the previously-tested frequencies, whereas in the previous report only about one quarter of the cells had rBMFs at the lowest tested frequency. Much of this disparity is likely due to testing at an extended range of MFs. Cells with rBMFs outside of the range of MFs previously tested make up about 40% of our sample, and many of these might not have registered sufficient activity in an AM search using the previous range of MFs to be further analyzed in the earlier experiment.

**Figure 3:**
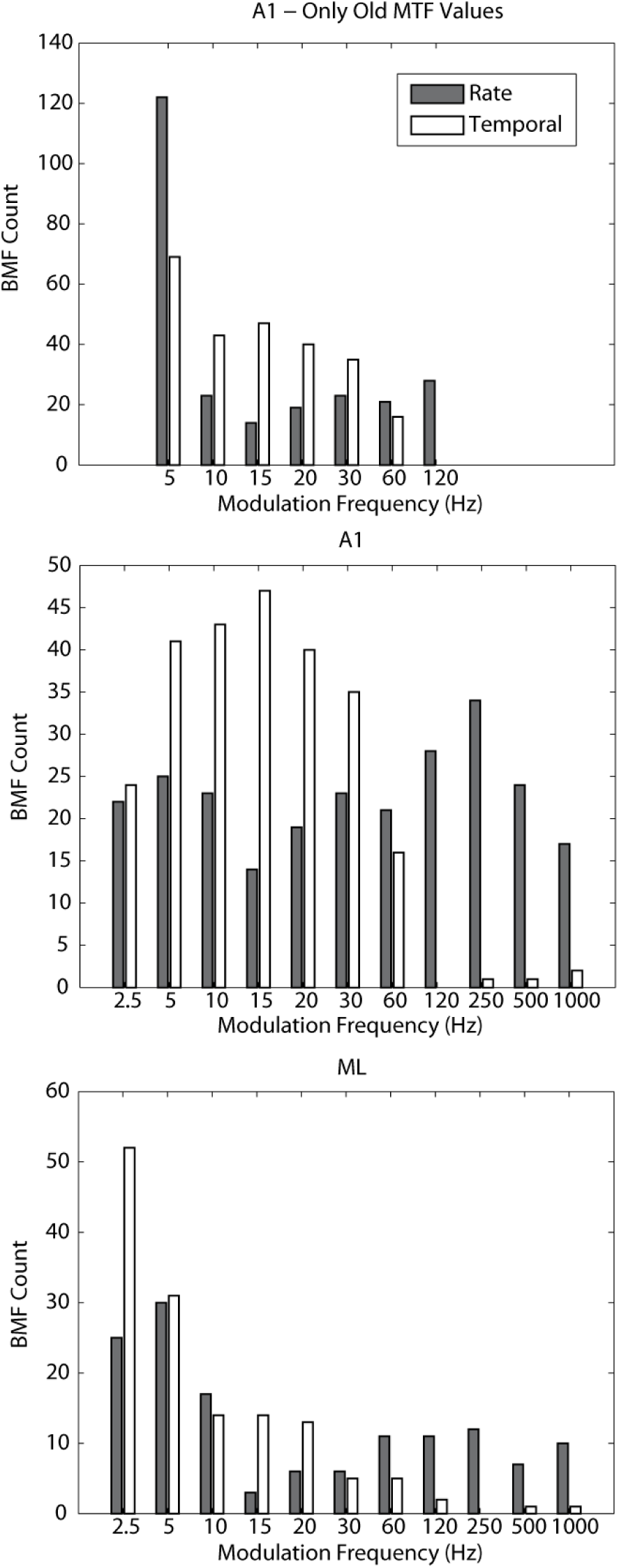
Best modulation frequencies. Top, best modulation frequencies (BMFs) for rate (shaded) and temporal (open) measures in A1 calculated using only the frequencies tested in a previous report (Yin et al. 2011). Middle, BMFs for A1 calculated using all frequencies tested in this study. Bottom, BMFs for ML calculated using all frequencies tested in this study.

Because of the wider range of modulation frequencies used for ML in this study it is important to compare these ML data to A1 data collected with the same range of modulation frequencies (i.e., the comparison to Yin et al. 2011 would not be a fair comparison). When looking at the fuller range of MFs, rBMF distributions in A1 are not significantly different from a flat distribution across modulation frequency (Kolmogorov-Smirnov test with counts adjusted for uneven sampling in log space, P = 0.07, ns), despite a small dip around 15 Hz (which may be expected as our sampling density temporarily increases on the log scale between 10 and 30 Hz) and a small peak around 250 Hz. tBMFs are clustered between 2.5 and 60 Hz as strong phase locking to higher modulation frequencies is rare in A1. In ML, both rBMFs and tBMFs show a shift towards lower frequencies relative to A1, with the vast majority of tBMFs at 20 Hz or below.

### Bandwidths (BWs)

For all bandpass fits (including E/S fits, discussed later) we calculated the associated bandwidths, defined as the full width of the fit at half-height. Some of our full widths extended below the lowest tested MF (2.5 Hz), and because we measure bandwidth in octaves this could occasionally result in very high-octave bandwidths where a large portion of the width lay outside of the tested MFs. In order to avoid this, we assigned a low frequency cutoff floor at 1.25 Hz and a high frequency cutoff ceiling of 2000 Hz (both one octave from the most extreme tested MFs) to ensure that the bandwidth values we calculated did not include excessive contributions from fits to untested MFs.

In general, we find a wide range of bandwidths. For A1 MTFs that were classified as bandpass (Fig. 4, upper left plot), the distribution of BWs is skewed towards lower widths, centered around 1-4 octaves for spike count (mean of 3.0 octaves) and a bit wider (centered around 1-6 octaves, mean of 3.7 octaves) for vector strength. Vector strength bandwidths are significantly wider than spike count (P = 6×10-4).

**Figure 4:**
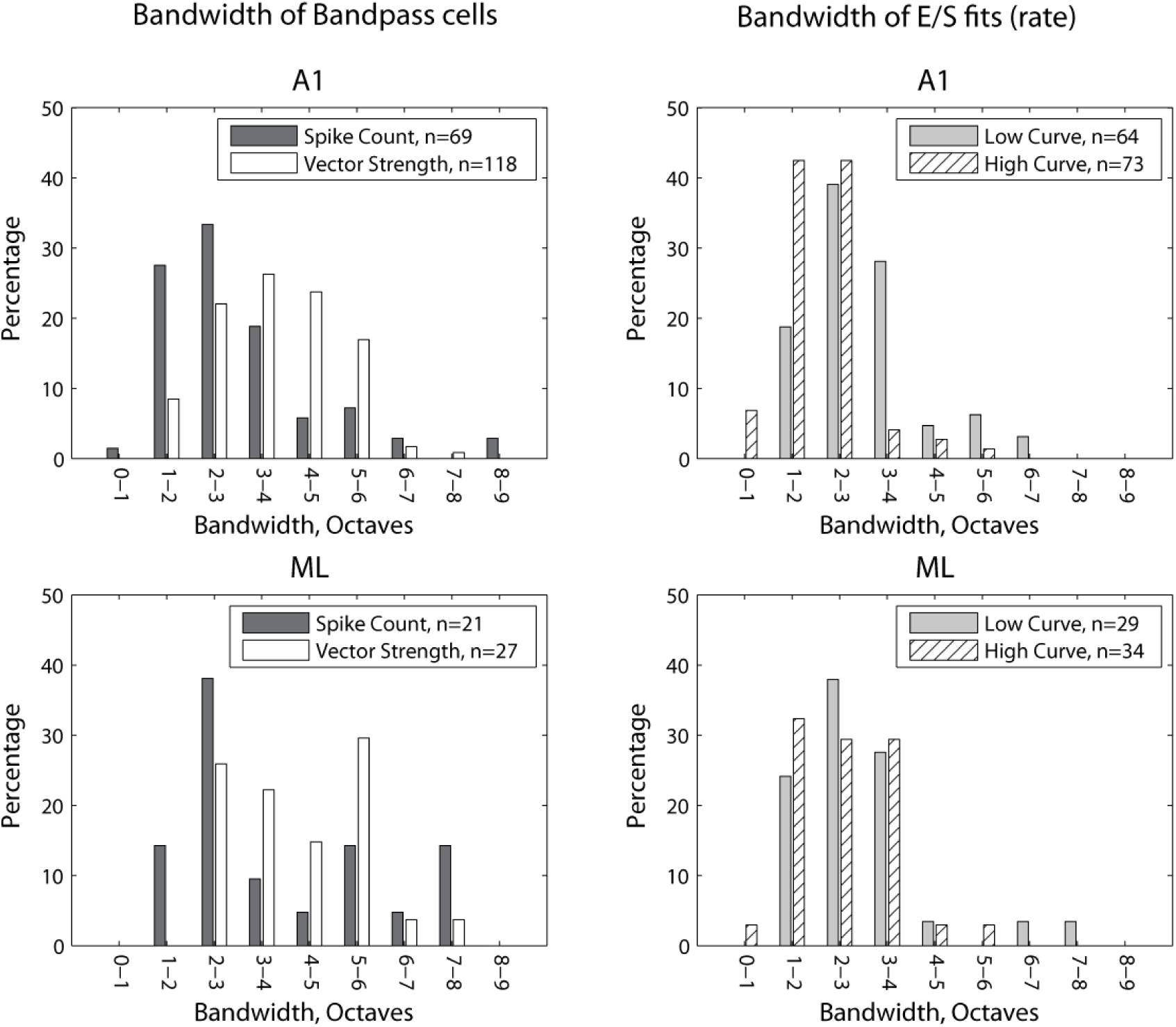
Distributions of bandwidths. Bandwidths are calculated as the octave width of the fitted curve at half-height (see Methods), constrained by upper and lower limits that are one octave outside of the range of tested modulation frequencies (MF). Top plots are A1 and bottom plots are ML. Left, bandwidths of bandpass cells for both rate and temporal measures. Right, rate bandwidths for the lower-MF peak (gray) and the higher-MF peak (hatched) for excitation/suppression cells. A small number of lower-MF peaks had low frequency cutoffs less than zero and were excluded from this analysis.

Bandwidth distributions in ML bandpass MTFs are not drastically different than those in A1. The mean spike count bandwidth is 3.8 octaves and the mean vector strength bandwidth is 4.1 octaves. Spike count and vector strength BW distributions did not differ significantly in ML (P = 0.53). Unlike A1, there are several cells with extremely large bandwidths. There is no significant difference between the bandwidth distributions in ML and A1 for either spike count (P = 0.06) or for vector strength (P = 0.13), though the trend in both cases is for wider fits in ML.

### Mean modulation transfer functions

To estimate the overall representation of different modulation frequencies, and any rolloff of the population of neurons at higher frequencies (Fig. 5) we calculated the mean modulation transfer functions. Psychophysically, macaque monkey modulation frequency sensitivity rolls off sharply between 250 and 1000 Hz (O’Connor et al. 2000; O’Connor et al. 2011). In A1, the mean rMTF of non-synchronized and of synchronized responses shows little MF dependence all the way to 1000 Hz (Solid lines in Fig, 5). In ML, it is difficult to determine whether the mean non-synchronized and synchronized rMTF are flat, but combining non-synchronized and synchronized flattens the curve even more.

**Figure 5:**
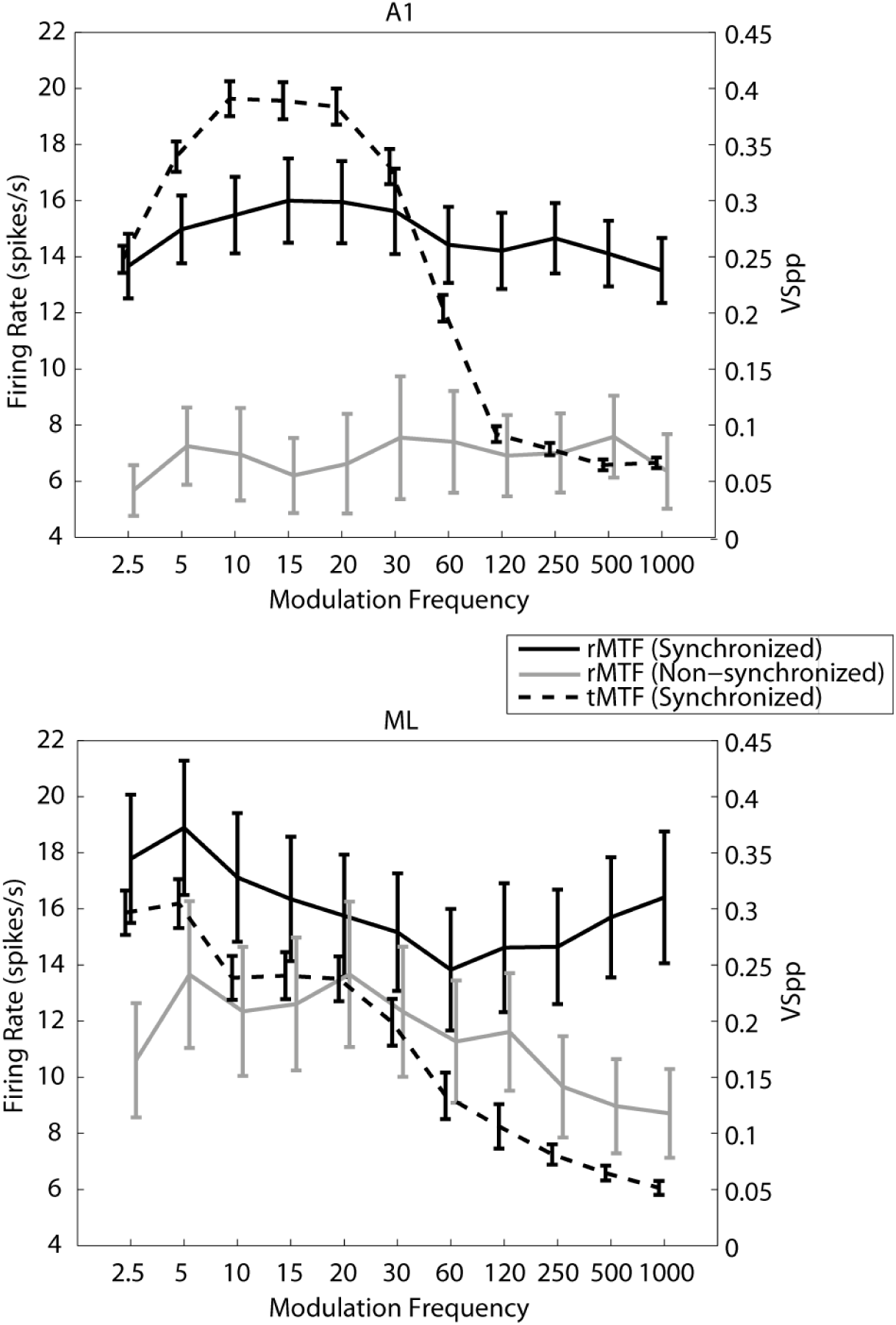
Mean modulation transfer functions (MTFs). Top plot is A1, bottom plot is ML. The mean MTF is calculated by taking the average response (firing rate or vector strength) of all cells at each MF. Only responsive cells are included. Rate modulation transfer functions (MTFs, solid lines) are referred to the left y-axis and broken down into synchronized and non-synchronized cells. Temporal MTFs (dotted lines) are referred to the right y-axis. Error bars indicate standard error of the mean. X-axis is slightly jittered to increase visibility of error bars.

The mean tMTF (dashed line) rolled off more steeply at high modulation frequencies than the mean rMTFs. A1 showed a sharp roll off beginning between 30-60 Hz, and the mean ML rMTF showed a more gradual roll off starting at 5 Hz.

### Temporal high frequency cutoffs

A common measure of phase-locking ability is the temporal high frequency cutoff – here defined as the highest modulation frequency at which a cell is able to significantly phase lock to an AM stimulus. We determined the temporal high frequency cutoff for both A1 and ML neurons, which is plotted in Fig. 6 as the percentage of recorded cells in each area with a cutoff at each tested MF. Because some cells do not significantly phase lock at all, the values in Fig. 6 do not add up to 100 percent. In addition to the reduction in number of cells that significantly phase lock in ML relative to A1 (see Table 1), there is also a downwards shift in the mean temporal cutoff in ML. To calculate the mean of the high frequency cutoffs, we used the geometric mean. Because MF is a non-linear octave-based measure, values to be averaged should first be transformed into an octave representation with the resulting mean transformed back into frequency, which is mathematically equivalent to taking the geometric mean of the frequency values. The mean A1 temporal high frequency cutoff is 27.7 Hz, significantly higher than the mean ML temporal high frequency cutoff of 19.5 Hz (two-tailed t-test, P = 0.034).

**Figure 6:**
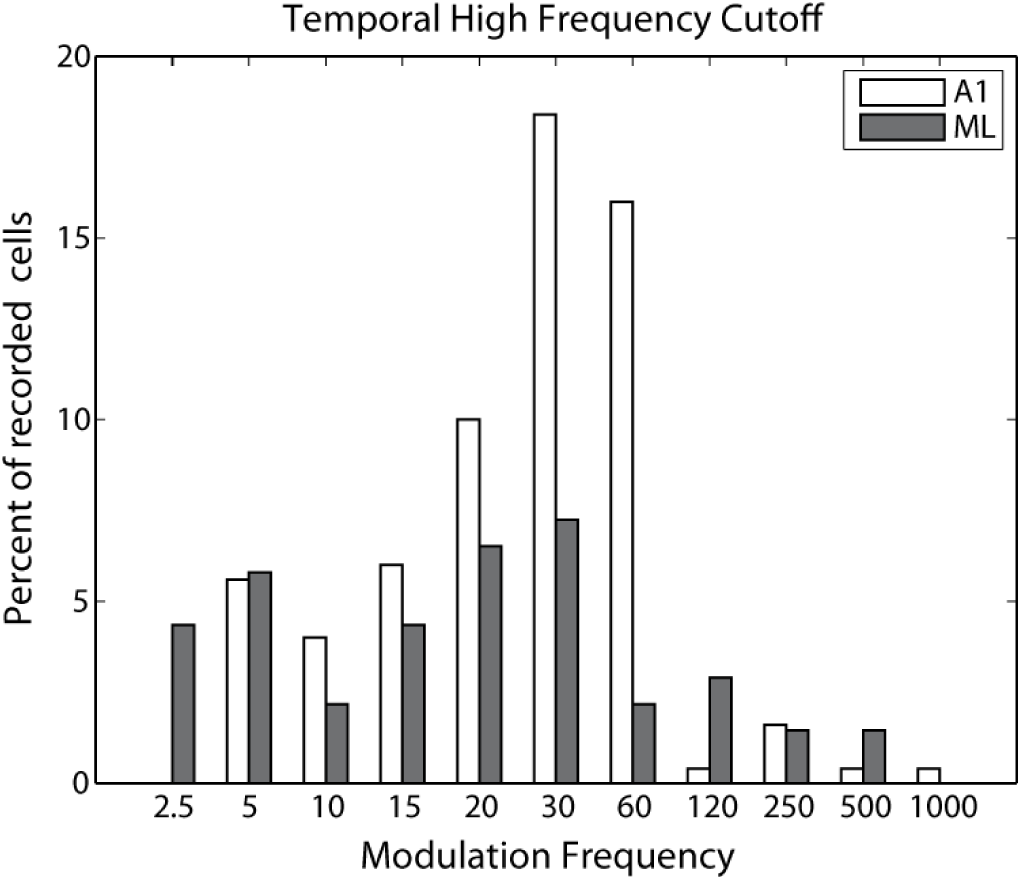
Temporal high frequency cutoffs. The temporal high frequency cutoff is defined as the highest modulation frequency which results in significant phase locking relative to an unmodulated stimulus. Percentages do not add up to 100 because some cells do not phase lock.

### Reliability of phase locking

In A1, it has been shown that at higher MFs, the reliability of phase locking – the likelihood that a neuron will fire synchronously on any given cycle – decreases even for cells with strong phase locking (Yin et al. 2011). Although some cells continue to fire in a temporally precise fashion to high MFs, they are unable to follow each AM cycle as the cycles get closer in time. Because reliability of phase locking drops off at higher MFs it may be related to a neuron’s high frequency temporal cutoff, which would suggest that ML neurons would show lower reliability than A1 neurons. In Fig. 7, we plot the average reliability of significantly phase-locking MFs for A1 and ML neurons recorded for this study (e.g. if for a given cell, it significantly phase locked to 15 Hz but not to 20 Hz, its 15 Hz responses would be included in the calculated mean, but its 20 Hz responses would not). Any MF with fewer than four significantly phase-locking cells was excluded from this analysis. Our measure of reliability uses cycle-by-cycle vector strength (*VS*_*CC*_, see Methods) and is calculated as *VS*_*CC*_/*VS*_*PP*_. This measure approximates the proportion of cycles that exhibit phase-locked firing. Fig. 7 shows that phase-locking ML neurons are not less reliable than phase-locking A1 neurons, despite the fact that the strength of phase locking (Fig. 5) and the high frequency cutoffs (Fig. 6) are lower in ML than in A1. Although the average phase-locking cell in ML has a response that is not as tightly phase locked as in A1, the reliability of that response is similar between the two areas.

**Figure 7:**
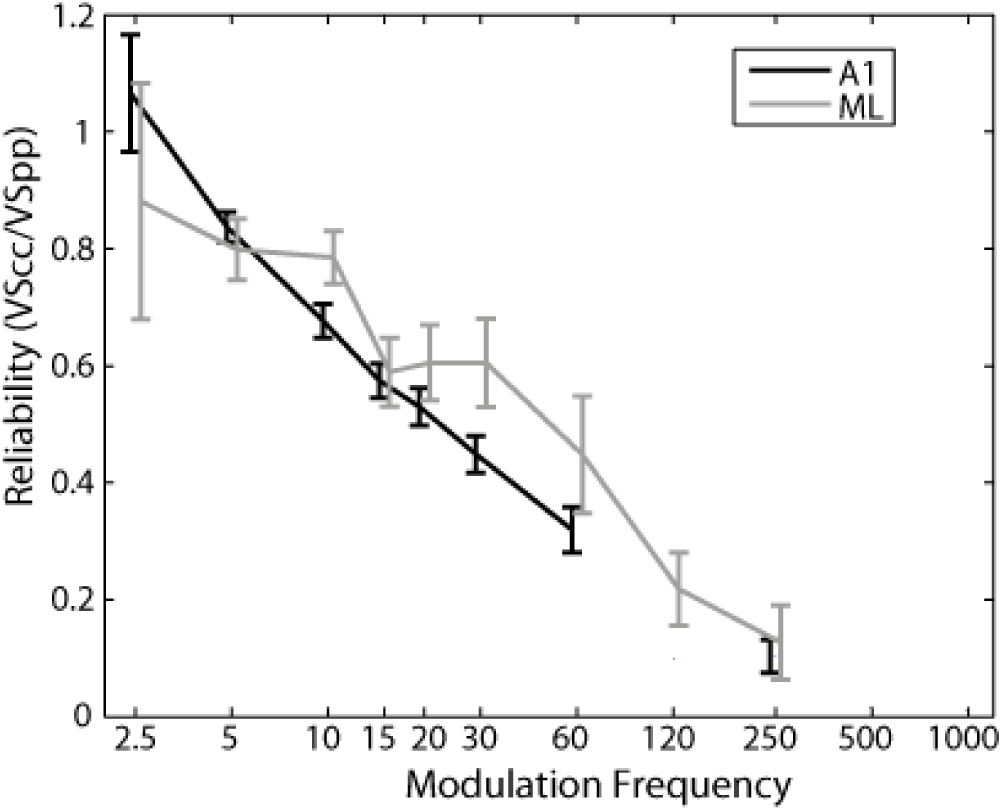
Reliability of synchronized firing. Mean reliability of synchronized firing in A1 and ML is estimated by the ratio of *VS*_*CC*_ to *VS*_*PP*_. Mean values are calculated only from modulation frequencies (MFs) that significantly phase lock; if fewer than 4 recorded cells significantly phase locked at a particular MF that point was omitted from the plot. Values above 1.0 (e.g. A1 at the 2.5 Hz MF) are due to fact that cycles that include the 70 ms onset response window are excluded from the *VS*_*CC*_ analysis but the corresponding times are not excluded from the *VS*_*PP*_ analysis. Plots are offset slightly left and right to allow better visibility of overlapping error bars. Error bars show standard error of the mean.

### Excitation/Suppression cells

As mentioned earlier, in both areas we find a class of cells that has previously not been widely reported in cortex – the E/S cells, which we only found for rate responses. These cells have two distinct regions of response – one region at low modulation frequencies which is usually synchronized (at a rate of about 80% in both A1 and ML, Table 1) and another region at high modulation frequencies which is rarely synchronized (about 5% in both A1 and ML, Table 1) separated by a region that is typically suppressed below the unmodulated response rate. In our previous report on A1 MTFs we were not able to effectively observe this class of cells because our tested MFs did not go high enough (only to 120 Hz, however note that the cell in Fig. 2A of Yin et al. 2011 also appears to be of this class but with a second response region that starts at lower MFs than commonly found). Because our tested MFs in the current experiment extended up to 1000 Hz, this class of cells became evident, and we developed new analyses (see Methods) to characterize them.

The E/S cells are found in high proportions in both A1 and ML, accounting for approximately 40% of all rate-responsive cells in each area. Figure 8 depicts four examples of this class of E/S cells, two from A1 and two from ML. The cells depicted in Figures 8A-C all show corresponding synchrony and firing rate in the lower response region, which is lost and then followed by a higher response region of non-synchronized firing at modulation frequencies of 250 Hz and above. The cell depicted in Figure 8D does not phase lock in either the lower frequency or higher frequency response region, which is less common for this type of cell.

**Figure 8:**
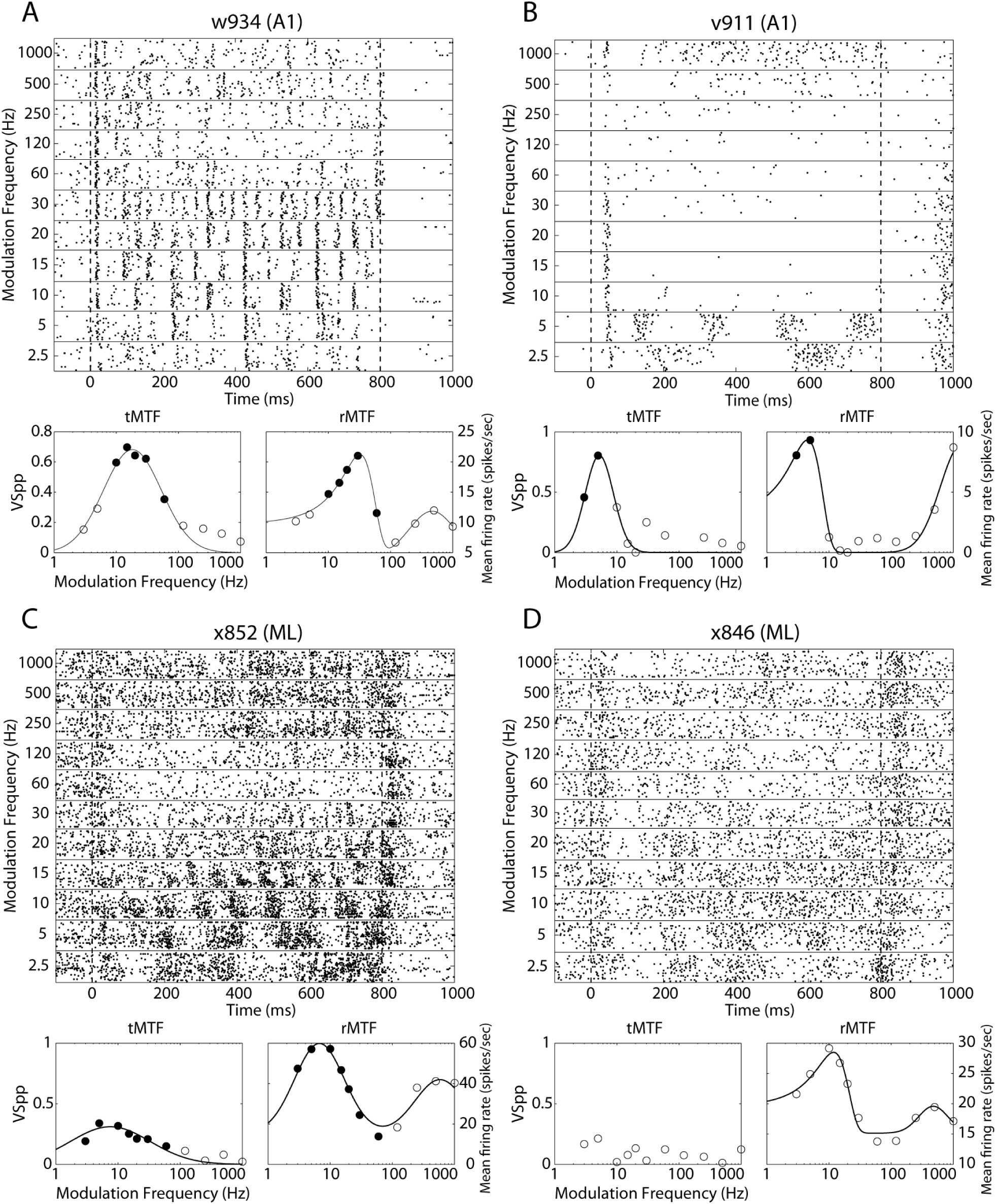
Example responses from excitation/suppression cells. Plots as in Figure 1. All examples are best fit by the excitation/suppression fit (see Methods). Filled dots indicate modulation frequencies at which there is significant phase locking (same modulation frequencies on both temporal and rate MTFs). Note that significant phase locking, when present, occurred exclusively within the low-MF peak of the excitation/suppression fit. The cell depicted in panel D does not significantly phase lock at any modulation frequency.

Examining a cell’s ability to detect modulation is particularly instructive with respect to firing rate measures in E/S cells. The firing rate at the lower-MF peak in E/S cells (Fig. 9, left) is almost exclusively higher than the unmodulated firing rate in both A1 and ML. At the same time, the firing rate at the suppressive trough does the opposite, being reduced to below the unmodulated firing rate (Fig. 9, middle) in both A1 and in ML. Although we have modeled the E/S cells as the sum of two positive peaks, the data is more suggestive of an excitatory peak flanked by an inhibitory region at higher MFs, followed by a return to the level of unmodulated firing at the highest MFs. Responses of the E/S cells to the highest tested MF of 1000 Hz (Fig. 9, right) are generally consistent with this. It appears that by and large neither A1 nor ML E/S cells are able to indicate the presence of high frequency (1000 Hz) AM by changing their firing rate but rather respond at the highest MFs as if to unmodulated noise. Given A1’s higher high frequency temporal cutoff relative to ML, the small number of significant differences in firing rate in A1 at a MF of 1000 Hz compared to almost none in ML is not surprising.

**Figure 9:**
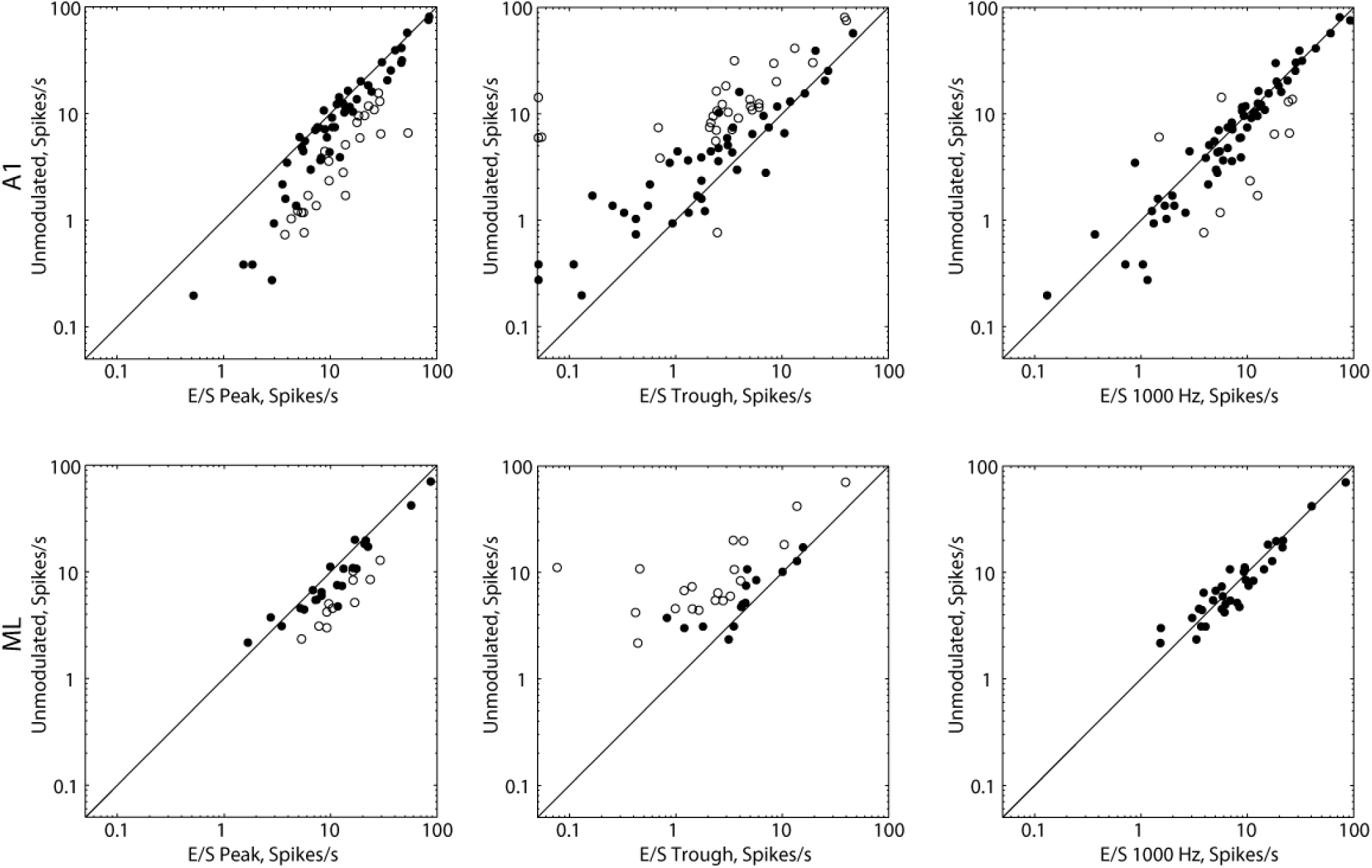
Firing rate relative to unmodulated noise at low frequency peak, band-reject trough, and high MF. Cell-by-cell scatter of unmodulated firing rate (y-axis) against various conditions. Left, firing rate of excitation/suppression (E/S) cells at the first E/S peak. Middle, firing rate of E/S cells at the E/S trough. Right, firing rate of E/S cells at the 1000 Hz modulation frequency (MF). Top row, A1. Bottom row, ML. Open symbols represent cells for which the two firing rates significantly differ (t-test, P < 0.05 after correction for multiple comparisons). Closed symbols represent cells for which firing rates do not significantly differ. Diagonal line is a unity line.

In comparison to the bandwidths of the purely bandpass cells, we also show the rate-based bandwidths of the cells which are categorized as E/S fits (Figure 4, right panels). In both A1 and ML the bandwidths of both the low-MF peak and the high-MF peak are as narrow or narrower than the rate bandwidth of the purely bandpass cells. For A1, the mean bandwidth of the low-MF peak is 3.0 octaves, the same value as seen for bandpass cells. The high-MF peak is significantly narrower than this at 2.1 octaves (P = 7×10-6).

In ML, the mean bandwidth of the low-MF peak is 2.9 octaves (compare to 3.8 octaves for BP cells in ML) while the mean bandwidth of the high-MF peak is 2.6 octaves – here the low-MF peak and the high-MF peak do not have significantly different bandwidths (P = 0.21). While the mean bandwidth of the low-MF peak does not change between A1 and ML (P = 0.85), the mean high-MF peak is wider in ML than in A1 (P = 0.04).

### Dual coding in AM-detecting cells

For cells that encode modulation by changing their spike rate, both increases and decreases in spike rate are observed. Most cells show exclusive increases or decreases in spike rate relative to unmodulated noise regardless of MF, while a smaller number of cells (“Mixed Inc/Dec”, commonly E/S cells) show increases in spike rate at some MFs but decreases at other MFs. Increases and decreases in spike rate can form a dual code – in isolation each coding scheme is potentially equally informative for detection, but a simple downstream summation of the activity of increasing and decreasing neurons would result in an impairment of detection.

Among cells that are exclusively non-synchronized (Table 2), an increasing code is more common than either a decreasing or mixed code in both A1 and ML (both comparisons of Increasing vs. Decreasing + Mixed Inc/Dec are significant at P < 0.01). Among synchronizing cells, the greater number of increasing cells in A1 looks very similar to that found for exclusively non-synchronizing cells (P < 0.001), but at the level of ML there is a shift to a nearly equal representation of increasing and decreasing cells (P = 0.69, ns), consistent with the findings of Niwa et al. 2013 in ML under active conditions. Thus, for synchronized cells, between the level of A1 and ML the ratio of increasing and decreasing cells in the population approaches a dual code, suggesting that this coding may provide an advantage in the detection of AM.

**Table 2:**
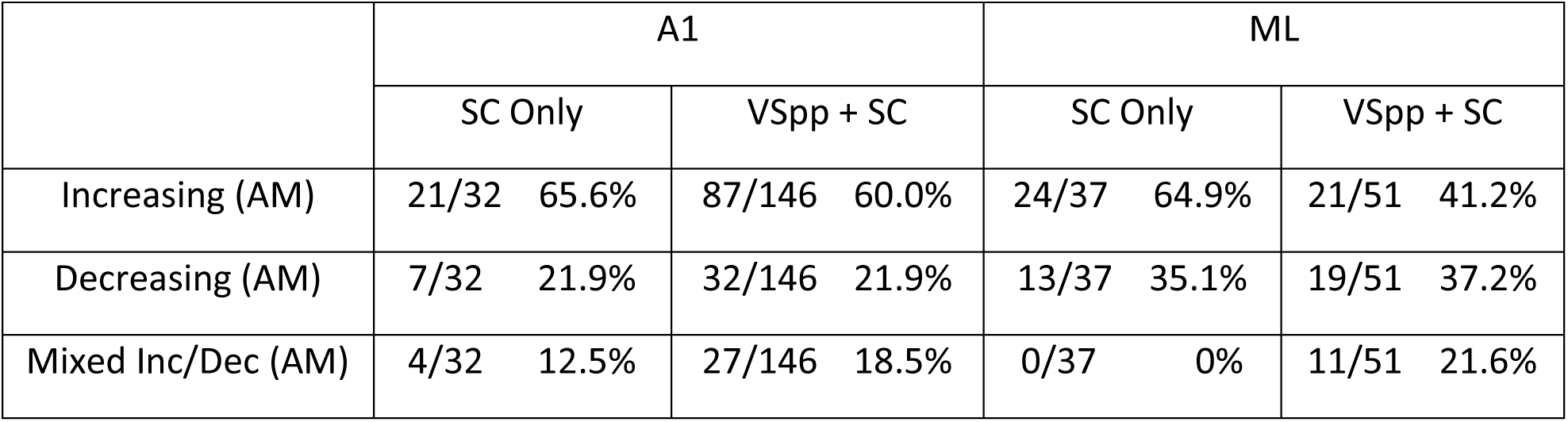
Dual Coding in AM-Detecting Cells. Values on the left are count of cells that detect AM by changing firing rate or phase locking, values on the right are in percentage. Cells in each area are broken down into synchronized (VSpp + SC) and exclusively non-synchronized (SC Only) categories. Detection of AM is also broken down into cells that show an increase or decrease in firing rate for AM relative to unmodulated, and “mixed” cells, which exhibit both increases and decreases in rate at different modulation frequencies.

## DISCUSSION

### AM processing in different monkey cortical fields

We compared basic neural response properties to AM noise in macaque cortical areas A1 and ML and found several similarities between the areas. The proportion of cells that encode AM by changing their firing rate is similar across the two areas, as are rMTF shapes. Response bandwidths are not statistically different across the two areas, though there is a slight trend for wider bandwidths in ML.

A1 and ML responses differ primarily in phase locking. We find several lines of evidence that suggest, in line with the typical temporal-to-rate transformation seen in the ascension of the auditory system, that phase locking is better in A1 than in ML – the proportion of cells that phase lock is higher in A1 (Table 1; Fig. 2); temporal BMFs are higher in A1 (Fig. 3); mean VSpp is higher in A1, particularly at lower MFs (Fig. 5); and high frequency phase-locking cutoffs are higher in A1 (Fig. 6). However, rate-based processing appears to be relatively similar in the two areas, suggesting that the proposed temporal-to-rate transformation with auditory system ascension (e.g. Schreiner and Urbas 1988; Lu et al. 2001; Lu and Wang 2004; Bendor and Wang 2007; Gao and Wehr 2015) may largely reflect a loss in temporal processing, rather than increasing activity for higher modulation frequencies.

A few studies have examined AM processing outside of A1 in monkeys. These have supported a nonstandard, nonhierarchical view of auditory cortex with regard to integration times and ability to phase lock. These studies suggest a possible integration time and corresponding phase-locking gradient in the caudomedial (short integration time, good phase locking) to anterolateral (long integration time, poor phase locking) direction; using a tone carrier, Scott et al. (2011) found that R, ML/AL (Anterior lateral belt of auditory cortex) and RM (rostromedial cortical field)/MM (middle medial cortical field) had similar phase-locking cutoff frequencies, which were lower than that of A1. The caudomedial cortical field, CM, however had phase locking similar to or better than A1. Since ML is both hierarchically higher than A1 and lateral, our data are consistent with both current hypothesis; that is, it might have lower frequency cutoffs and longer integration times because it is hierarchically higher than A1 or because it is more lateral in the caudomedial-to-anterolateral direction.

It is worthwhile to note that the rostrolateral-to-caudomedial axis has also been suggested as a possible gradient in the influence of somatosensory information in A1 with the caudomedial direction having the strongest somatosensory influence (Fu et al. 2003), as well as a caudal to rostral axis for spatial (caudal) versus non-spatial (rostral) processing (Romanski et al. 1999; Rauschecker and Tian 2000; Camalier et al. 2012). The degree to which these functional axes interact to create a functional organization of the auditory cortex has not yet been resolved.

The A1 population mean MTF data (Fig. 5) is qualitatively similar to that obtained by Scott et al. (2011), who measure the percentages of cells with significant firing rate or phase locking at each modulation frequency for tone-carrier sine-AM in both A1 and the core field R. They found a flat population rate function and a temporal MTF with a sharp rolloff in A1. However, the slope of the rolloff in A1 is slightly shallower in Scott et al. (2011) than what we observed. R also had a flat population rMTF, but R’s temporal MTF had a much sharper rolloff than what we observe in ML, consistent with R being in the core of auditory cortex and ML in the belt. These results are consistent in showing that the cortical representation of AM can neither be fully accounted for by rate responses alone nor by temporal responses alone – both studies suggest that the behavioral cutoff frequency for detection of AM (∼250-500 Hz) is higher than the phase-locking cutoff in cortical cells. The presence of relatively flat population rate MTFs suggests that rate coding can contribute to AM perception at higher modulation frequencies, but overall sensitivity may be lower because of a reduced contribution from phase-locked responses.

### Species and stimulus effects

AM has been studied in many species (reviewed in Joris et al. 2004). Across a wide range of measures, such as MTF tuning, temporal following as the auditory hierarchy is ascended, sensitivity of temporally phase-locked versus rate codes, and emergence of rate codes with auditory system ascension, qualitative results are similar across species (e.g. Liang et al. 2002; Nelson and Carney 2007; Ter-Mikaelian et al. 2007; Rosen et al. 2010; Hoglen et al. 2018). However, there are many specific quantitative differences in these measures. Macaques, including those in this study, fall somewhere between rodents (e.g. Ter-Mikaelian et al. 2007; Rosen et al. 2010; Hoglen et al. 2018) with reduced neural AM frequency following and sensitivity and new world monkeys (e.g., Liang et al. 2002; Hoglen et al 2018;) with high sensitivity and envelope following.

The type of acoustic stimulus used has particular significance when studying temporal processing. We used sinusoidal AM-noise. Most other studies of temporal processing have used different carriers, as well as click trains. Malone et al. (2013) showed that the carrier could have large effects on single neuron responses, but little effect on population statistics. Results with clicks are more problematic to interpret in relation to AM. Clicks are fundamentally different stimuli, even though they share a broad-band power spectrum with our AM noise stimuli. Click trains comprise repeated discrete pulses whose power is restricted to (more or less) precise times. In contrast, AM combines gradual temporal envelope fluctuations and periodicity, and it is well established that auditory cortical neurons are highly sensitive to envelope properties (Heil 2003). Click trains tend to yield a higher proportion of non-synchronized responses at fast click rates than is observed with AM (e.g. Lu et al. 2001; Liang et al. 2002; Yin et al. 2011; Gao and Wehr 2015). This may be a result of limited temporal resolution (similar to flicker fusion in vision, or what we see at higher modulation rates). There have been studies attempting to bridge the gap between clicks and AM by using stimuli constructed to be intermediate between amplitude modulation and click trains by varying duty cycle and envelope (e.g., Krebs et al. 2008; Lee et al. 2016; Osman et al. 2017). Lee et al. show a loss of temporal precision for high rates/envelopes, but they did not address whether this was due to a concomitantly improved rate code or simply a loss of fidelity. An intracellular study with click trains (Gao and Wehr 2015) has shown that fast click trains in rat AC cause intracellular depolarized plateaus (∼20 mv) with very low voltage click following (∼0.2 mV). By comparing currents under voltage clamp to voltage changes over time, the authors conclude that low pass filtering by membrane capacitance cannot account for the non-synchronized responses at high click repetition rates. This suggests there is some specialized mechanism for processing rapid discrete events. However, these results are difficult to compare to ours since there is no analog to unmodulated carriers for click trains, and mechanisms other than membrane capacitance could make it impossible for cells to temporally follow rapid discrete events with fast rise time.

### Dual coding of AM

The idea of a ‘dual’ opponent code for AM – one in which cells can code AM using either increases or decreases in firing rate – has a long history. Krishna and Semple (2000) found this effect in the inferior colliculus of gerbils when presenting AM tones, and Salinas et al. (2000) found a related frequency-dependent effect in somatosensory cortex of macaques when presenting them with fluttering vibrotactile stimuli. Yin et al. 2011 also reported finding cells in macaque A1 showing decreases in firing rate (relative to unmodulated) to detect AM. Here we find significantly more increasing cells relative to decreasing cells in A1 than found in Yin et al. 2011. This discrepancy is a bit difficult to explain, but could be due to differences in the level of behavioral training in the animals. In the present study highly trained animals alternated between passive blocks (no task) and blocks where they performed an AM detection task. We found that increasing cells outnumber decreasing cells by about three-to-one in A1, but that there is a shift towards decreasing cells in ML, where we see a slightly less than two-to-one ratio for non-synchronized cells and a near one-to-one ratio for synchronized cells. Nearly identical proportions of increasing and decreasing responses were found for A1 and ML neurons in these same animals while they were performing the AM detection task (Niwa et al. 2013), suggesting this is not an effect of task engagement.

### Excitation/Suppression cells and faux AM sensitivity

We found that a large percentage of cells have a second response region at higher modulation frequencies (Fig. 8) which almost usually was indistinguishable from the response to unmodulated noise (Fig. 9, right panels). This is likely because increasingly rapid modulation envelopes are difficult for neurons to follow (and perceptually difficult to detect). This finding suggests that few cells in A1 and almost no cells in ML can detect AM at very high frequencies, and instead they act as if they are responding to the noise carrier.

This point is particularly important because very few physiological studies of AM processing compare responses to the unmodulated carrier. Therefore, reports of non-synchronized responses to AM in auditory cortex are likely overestimated because many of those responses might simply be responses to the carrier, with no contribution from the modulation. It seems that any study interpreting non-synchronized activity at high modulation frequencies should make sure to rule out the possibility that rather than responding to modulation, the neuron is responding solely to the carrier.

We have used the term “excitation/suppression” (E/S) to describe these cells rather than the more commonly used ‘multi-peak’ or ‘band-reject’. “Multi-peak” suggests that there are two regions of response to modulation (it is unclear in most cases whether the second peak contains any response to the modulation as opposed to response to the carrier) and “band-reject” gives the impression that the primary response is a suppression, while not acknowledging the excitatory peak at lower modulation frequencies.

Cells with MTFs similar to our E/S cells have been previously characterized for AM tones in the inferior colliculus of rabbit (Nelson and Carney 2007) and gerbil (Krishna and Semple 2000; Krebs et al. 2008), for AM noise in the inferior colliculus of cat (Zheng and Escabí 2008), and for AM using tone and noise carriers in auditory thalamus of rat (Cai and Caspary 2015). Examples of this type of cell (often called “band-reject”) are not so widely reported in cortex. Outside of Bieser and Müller-Preuss (1996), who reported that about 20% of cells in squirrel monkey auditory cortex were “multipeak”, most examples of these cells in primate auditory cortex found in the literature have gone mainly unremarked upon (e.g. for AM tones in Barbour and Wang 2002; Liang et al. 2002; Lu and Wang 2004; for AM noise in Yin et al. 2011), perhaps because they were not prevalent enough to constitute a clearly-defined category. Here we find that in both A1 and ML, about 40% of cells exhibit an E/S MTF, likely due to our expanded modulation frequency test range. This value of 40% is similar to the value found in auditory thalamus (Cai and Caspary, 2015), which is suggestive that the E/S property may be inherited from lower levels of the auditory hierarchy, but the possibility that this property might be recalculated *de novo* in cortex cannot be discounted.

We have fit our E/S cells, which generally appear to have two positive-going peaks, with two positive Gaussians. However, considering the fact that the entire population (both A1 and ML, for all MTF types) appears to be similar to the unmodulated response at very high modulation frequencies, this may not be the most appropriate way to describe the cells. As our terminology (“excitation/suppression”) suggests, there appears to be a lower frequency positive peak flanked by a middle-frequency suppressive region, where responses typically fall below the response to the unmodulated, followed by a recovery towards the unmodulated “baseline”. At the same time, many neurons (e.g. Fig. 1B, Fig. 9A, Fig. 9D) do appear to have a downturn at 1000 Hz, suggesting that some excitatory drive may exist in the region of the second peak – whether this is a second region of excitatory response to modulation or simply the remnants of a very wide lower peak emerging from a narrower suppression is not clear. We note that in A1, the average bandwidths of the BP cells and the lower band of the E/S cells are essentially the same, which might argue against the higher peak being the tail of a very wideband response emerging from a narrower suppression. On the other hand, we do see very wideband BP responses (e.g. Fig. 1D) so this is not out of the question. Unlike in A1, in ML the average bandwidth of the lower band of the E/S cells is narrower than the bandwidth of the BP cells, which is suggestive of flanking inhibition sharpening the tuning of the E/S cells.

## Acknowledgements

This work was funded by NIH NIDCD grant DC02514 (MLS)

## REFERENCES

Barbour DL, Wang X. Temporal coherence sensitivity in auditory cortex. J Neurophysiol 88: 2684–2699, 2002.

Bendor D, Wang X. Differential neural coding of acoustic flutter within primary auditory cortex. Nat Neurosci 10: 763–771, 2007.

Bendor D, Wang X. Neural response properties of primary, rostral, and rostrotemporal core fields in the auditory cortex of marmoset monkeys. J Neurophysiol 100: 888–906, 2008.

Bieser A, Müeller-Preuss P. Auditory responsive cortex in the squirrel monkey: neural responses to amplitude-modulated sounds. Exp Brain Res 108:273–284, 1996.

Bohlen P, Dylla M, Timms C, Ramachandran R. Detection of modulated tones in modulated noise by non-human primates. J Assoc Res Otolaryngol 15: 801–821, 2014.

Bregman AS. Auditory Scene Analysis. Cambridge, MA: MIT Press, 1990.

Cai R, Caspary DM. GABAergic inhibition shapes SAM responses in rat auditory thalamus. Neurosci 299:146–155, 2015.

Camalier CR, D’Angelo WR, Sterbing-D’Angelo SJ, de la Mothe LA, Hackett TA. Neural latencies across auditory cortex of macaque support a dorsal stream supramodal timing advantage in primates. Proc Natl Acad Sci 109: 18168–18173, 2012.

Cohen YE, Theunissen F, Russ BE, Gill P. Acoustic features of rhesus vocalizations and their representation in the ventrolateral prefrontal cortex. J Neurophysiol 97: 1470–1484, 2007.

Drullman R, Festen JM, Plomp R. Effect of temporal envelope smearing on speech reception. J Acoust Soc Am 95: 2670–2680, 1994.

Elliott TM, Theunissen FE. The modulation transfer function for speech intelligibility. PLoS Comput Biol (March 6, 2009). doi: 10.1371/journal.pcbi.1000302.

Fu KM, Johnston TA, Shah AS, Arnold L, Smiley J, Hackett TA, Garraghty PE, Schroeder CE. Auditory cortical neurons respond to somatosensory stimulation. J Neurosci 23:7510–7515, 2003.

Gao X, Wehr M. A coding transformation for temporally structured sounds within auditory cortical neurons. Neuron 86: 292–303, 2015.

Geffen MN, Gervain J, Werker JF, Magnasco MO. Auditory perception of self-similarity in water sounds. Front Integr Neurosci (May 11, 2011). doi: 10.3389/fnint.2011.00015.

Gervain J, Geffen MN. Efficient neural coding in auditory and speech perception. Trends Neurosci 42: 56–65, 2019.

Goldberg JM, Brown PB. Response of binaural neurons of dog superior olivary complex to dichotic tonal stimuli: some physiological mechanisms of sound localization. J Neurophysiol 32: 613–636, 1969.

Grimault N, Bacon SP, Micheyl C. Auditory stream segregation on the basis of amplitude-modulation rate. J Acoust Soc Am 111: 1340–1348, 2002.

Heil P. Coding of temporal onset envelope in the auditory system. Speech Comm 41: 123–134, 2003.

Hershenhoren I, Nelken I. Detection of tones masked by fluctuating noise in rat auditory cortex. Cereb Cortex 27: 5130–5143, 2017.

Hoglen NEG, Larimer P, Phillips EAK, Malone BJ, Hasenstaub AR. Amplitude modulation coding in awake mice and squirrel monkeys. J Neurophysiol 119: 1753–1766, 2018.

Itatani N, Klump GM. Auditory streaming of amplitude-modulated sounds in the songbird forebrain. J Neurophysiol 101: 3212–3225, 2009.

Jin SH, Nelson PB. Speech perception in gated noise: the effects of temporal resolution. J Acoust Soc Am 119: 3097–3108, 2006.

Johnson JS, Yin P, O’Connor KN, Sutter ML. Ability of primary auditory cortical neurons to detect amplitude modulation with rate and temporal codes: neurometric analysis. J Neurophysiol 107:3325–3341, 2012.

Joris PX, Schreiner CE, Rees A. Neural processing of amplitude-modulated sounds. Physiol Rev 84: 541–577, 2004.

Kaas JH, Hackett TA. Subdivisions of auditory cortex and processing streams in primates. Prod Natl Acad Sci 97:11793–11799, 2000.

Krebs B, Lesica NA, Grothe B. The representation of amplitude modulations in the mammalian auditory midbrain. J Neurophysiol 100: 1602–1609, 2008.

Krishna BS, Semple MN. Auditory temporal processing: responses to sinusoidally amplitude-modulated tones in the inferior colliculus. J Neurophysiol 84: 255–273, 2000.

Lee CM, Osman AF, Volgushev M, Escabí MA, Read HL. Neural spike-timing patterns vary with sound shape and periodicity in three auditory cortical fields. J Neurophysiol 155: 1886–1904, 2016.

Liang L, Lu T, Wang X. Neural representations of sinusoidal amplitude and frequency modulations in the primary auditory cortex of awake primates. J Neurophysiol 87: 2237–2261, 2002.

Liu RC, Miller KD, Merzenich MM, Schreiner CE. Acoustic variability and distinguishability among mouse ultrasound vocalizations. J Acoust Soc Am 114: 3412–3422, 2003.

Lu T, Liang L, Wang X. Temporal and rate representations on time-varying signals in the auditory cortex of awake primates. Nat Neurosci 4: 1131–1138, 2001.

Lu T, Wang X. Information content of auditory cortical responses to time-varying acoustic stimuli. J Neurophysiol 91: 301–313, 2004.

Malone BJ, Scott BH, Semple MN. Dynamic amplitude coding in the auditory cortex of awake rhesus macaques. J Neurophysiol 98: 1451–1474, 2007.

Malone BJ, Beitel RE, Vollmer M, Heiser MA, Schreiner CE. Spectral context affects temporal processing in awake auditory cortex. J Neurosci 33: 9431–9450, 2013.

Mardia KV, Jupp PE. Directional Statistics. New York: Wiley, 2000.

Narayan R, Graña G, Sen K. Distinct time scales in cortical discrimination of natural sounds in songbirds. J Neurophysiol 96: 252–258, 2006.

Nelson PC, Carney LH. Neural rate and timing cues for detection and discrimination of amplitude-modulated tones in the awake rabbit inferior colliculus. J Neurophysiol 97: 522–539, 2007.

Niwa M. Johnson JS, O’Connor KN, Sutter ML. Active engagement improves primary auditory cortical neurons’ ability to discriminate temporal modulation. J Neurosci 32: 9323–9334, 2012.

Niwa M, Johnson JS, O’Connor KN, Sutter ML. Differences between primary auditory cortex and auditory belt related to encoding and choice for AM sounds. J Neurosci 33: 8378–8395, 2013.

Niwa M, O’Connor KN, Engall E, Johnson JS, Sutter ML. Hierarchical effect of task engagement on amplitude modulation encoding in auditory cortex. J Neurophysiol 113: 307–327, 2015.

O’Connor KN, Barruel P, Sutter ML. Global processing of spectrally complex sounds in macaques (*Macaca mullata*) and humans. J Comp Physiol 186: 903–912, 2000.

O’Connor KN, Johnson JS, Niwa M, Noriega NC, Marshall EA, Sutter ML. Amplitude modulation detection as a function of modulation frequency and stimulus duration: comparisons between macaques and humans. Hear Res 277: 37–43, 2011

Osman AF, Lee, CM, Escabí MA, Read HL. A hierarchy of time scales for discriminating and classifying the temporal shape of sound in three auditory cortical fields. J Neurosci 38: 6967–6982, 2018.

Overton JA, Recanzone GH. Effects of aging on the response of single neurons to amplitude-modulated noise in primary auditory cortex of rhesus macaque. J Neurophysiol 115: 2911–2923, 2016.

Rauschecker JP, Tian B. Mechanisms and streams for processing of “what” and “where” in auditory cortex. Proc Natl Acad Sci 97: 11800–11806, 2000.

Rauschecker JP, Tian B. Processing of band-passed noise in the lateral auditory belt cortex of the rhesus monkey. J Neurophysiol 91:2578–2589, 2004.

Rauschecker JP, Tian B, Hauser M. Processing of complex sounds in the macaque nonprimary auditory cortex. Science 268:111–114, 1995.

Romanski LM, Tian B, Fritz J, Mishkin M, Goldman-Rakic PS, Rauschecker JP. Dual streams of auditory afferents target multiple domains in the primate prefrontal cortex. Nat Neurosci 2: 1131–1136, 1999.

Rosen MJ, Semple MN, Sanes DH. Exploiting development to evaluate auditory encoding of amplitude modulation. J Neurosci 30: 15509–15520, 2010.

Rosen S. Temporal information in speech: acoustic, auditory and linguistic aspects. Philos Trans R Soc Lond B Biol Sci 336: 367–373, 1992.

Salinas E, Hernández A, Zainos A, Romo R. Periodicity and firing rate as candidate neural codes for the frequency of vibrotactile stimuli. J Neurosci 20: 5503–5515, 2000.

Schreiner CE, Urbas JV. Representation of amplitude modulation in the auditory cortex of the cat. II. Comparison between cortical fields. Hear Res 32: 49–64, 1988.

Scott BH, Malone BJ, Semple MN. Transformation of temporal processing across auditory cortex of awake macaques. J Neurophysiol 105: 712–730, 2011.

Shannon RV, Zeng FG, Kamath V, Wygonski J, Ekelid M. Speech recognition with primarily temporal cues. Science 270: 303–304, 1995.

Singh NC, Theunissen FE. Modulation spectra of natural sounds and ethological theories of auditory processing. J Acoust Soc Am 114: 3394–3411, 2003.

Steinschneider M, Fishman YI, Arezzo JC. Representation of the voice onset time (VOT) speech parameter in population responses within primary auditory cortex of the awake monkey. J Acoust Soc Am 114: 307–321, 2003.

Ter-Mikaelian M, Sanes DH, Semple MN. Transformation of temporal properties between auditory midbrain and cortex in the awake Mongolian gerbil. J Neurosci 27: 6091–6102, 2007.

Tian B, Rauschecker JP. Processing of frequency-modulated sounds in the lateral auditory belt cortex of the rhesus monkey. J Neurophysiol 92: 2993–3013, 2004.

Tian B, Reser D, Durham A, Kustov A, Rauschecker JP. Functional specialization in rhesus monkey auditory cortex. Science 292:290–293, 2001.

Woods TM, Lopez SE, Long JH, Rahman JE, Recanzone GH. Effects of stimulus azimuth and intensity in the single-neuron activity in the auditory cortex of the alert macaque monkey. J Neurophysiol 96: 3323–3337, 2006.

Xiang J, Poeppel D, Simon JZ. Physiological evidence for auditory modulation filterbanks: cortical responses to concurrent modulations. J Acoust Soc Am 133:EL7–EL12, 2013.

Yamagishi S, Otsuka S, Furukawa S, Kashino M. Comparison of perceptual properties of auditory streaming between spectral and amplitude modulation domains. Hear Res 350: 244–250, 2017.

Yin P, Johnson JS, O’Connor KN, Sutter ML. Coding of amplitude modulation in primary auditory cortex. J Neurophysiol 105: 582–600, 2011.

Yost WA. Auditory image perception and analysis: the basis for hearing. Hear Res 56: 8–18, 1991.

Zeng FG, Nie K, Stickney GS, Kong YY, Vongphoe M, Bhargave A, Wei C, Cao K. Speech recognition with amplitude and frequency modulations. Proc Natl Acad Sci USA 102: 2293–2298, 2005.

Zheng Y, Escabí MA. Distinct roles for onset and sustained activity in the neuronal code for temporal periodicity and acoustic envelope shape. J Neurosci 28: 14230–14244, 2008.

